# MaGIC: a machine learning tool set and web application for monoallelic gene inference from chromatin

**DOI:** 10.1101/353359

**Authors:** Svetlana Vinogradova, Sachit D. Saksena, Henry N. Ward, Sébastien Vigneau, Alexander A. Gimelbrant

**Affiliations:** Department of Cancer Biology, Dana-Farber Cancer Institute, Boston, MA 02115; Department of Genetics, Harvard Medical School, Boston, MA 02115; Institute for Cellular and Molecular Biology, Department of Molecular Biosciences, The University of Texas at Austin, Austin, TX 78712; University of Minnesota-Twin Cities, Bioinformatics and Computational Biology Program, Minneapolis, MN 55455

## Abstract

**Summary:** A large fraction of human and mouse autosomal genes are subject to random monoallelic expression (MAE), an epigenetic mechanism characterized by allele-specific gene expression that varies between clonal cell lineages. MAE is highly cell-type specific, and mapping it in a large number of cell and tissue types can provide insight into its biological function. Its detection, however, remains challenging. We previously reported that a sequence-independent chromatin signature identifies, with high sensitivity and specificity, genes subject to MAE in multiple tissue types using readily available ChIP-seq data. Here we present an implementation of this method as a user-friendly, open-source software pipeline for **<u>m</u>**ono**<u>a</u>**llelic **<u>g</u>**ene **<u>i</u>**nference from **<u>c</u>**hromatin (MaGIC).

**Availability and implementation:** The source code for the MaGIC pipeline and the Shiny app is available at https://github.com/gimelbrantlab/magic.

## Introduction

Autosomal random monoallelic expression (MAE), like X-chromosome inactivation and imprinting, is a mitotically stable epigenetic mechanism that profoundly affects genotype-phenotype relationship in mammals by controlling the relative expression of the two parental alleles (Chess, 2016; Savova *et al.*, 2013). MAE is the most widespread of these three phenomena, affecting over 10% of human autosomal genes, including multiple genes implicated in cancer, autism, and Alzheimer’s disease. Similar to X-inactivation, MAE is clone-specific and thus is not detected in polyclonal, tissue-level measurement of allelic imbalance in transcription. As an alternative to direct measurement, we have identified a chromatin signature of monoallelic expression, which can be applied to detect MAE in polyclonal samples (Nag *et al.*, 2013).

This chromatin signature is based on gene-body enrichment of histone H3 Lys-27 trimethylation (H3K27me3) and H3 Lys-36 trimethylation (H3K36me3) as measured by ChIP-seq. We experimentally confirmed the signature’s accuracy in multiple human and mouse cell-types using clonal cell lines with known allelic expression as a reference (Nag *et al.*, 2013, Nag *et al.*, 2015). Use of the chromatin-based MAE maps has already led to new insights in genome evolution (Savova *et al.*, 2016) and neurodevelopmental disease (Savova *et al.*, 2017). However, to enable a broader adoption of this approach on a variety of large-scale datasets, an integrated software tool was needed.

Here, we describe a user-friendly, flexible, open-source software pipeline named **<u>m</u>**ono**<u>a</u>**llelic **<u>g</u>**ene **<u>i</u>**nference from **<u>c</u>**hromatin (MaGIC). In addition to classifying genes as MAE or biallelic based on existing models, MaGIC can also generate predictive models *de novo*, as new, larger datasets become available.

### The MaGIC Pipeline

MaGIC is a command-line software package written in R that consists of three separate parts (**Figure 1a**).

**Process.R** processes ChIP-seq bigWig (Kent *et al.*, 2010) files into gene-body or promoter enrichment normalized to control data (e.g. input or whole-cell extract) or to feature length. First, Bwtool (Pohl and Beato, 2014) is used to calculate mean signals for gene intervals based on a reference annotation. X-linked, imprinted, and olfactory receptor genes are then filtered out by default to focus on the less characterized autosomal random MAE genes. Intervals with control signals lower than a user-defined threshold are also removed. Finally, ChIP-seq signals are normalized to control signals, and the resulting values are converted to quantile rank and saved to a file. This output file can be used to generate new classifiers using *generate.R* or to predict monoallelic expression using *analyze.R* with pre-trained classifiers.

**Generate.R** trains classifiers on ChIP-seq data processed by the *process.R*, script using a variety of algorithms supported by the *caret* R package (Kuhn, 2008). A set of training labels containing true allelic expression calls for genes, typically determined by RNA-Seq in related clonal cell lines, should be provided in a separate file. Alternatively, the user can utilize one of the training label sets we provide, which corresponds to human or mouse B lymphoid cells. By default, *generate.R*, trains models with five-fold crossvalidation, although the degree of cross-validation can be modified by the user. If the user has a separate validation set, they can instead train the classifier on the complete set of training data. In all cases, *generate.R* outputs a set of models and a summary file containing per-model performance metrics.

**Analyze.R** predicts monoallelic expression using ChIP-seq data processed by *process.R* and classifiers generated by *generate.R*. The predictions can be filtered by gene length and expression level, provided by the user in a separate file, with default threshold values as in (Nag *et al.*, 2015). The output from *analyze.R* contains predicted allelic expression status by gene.

**Shiny web application:** To make MaGIC more user-friendly and add additional visualizations, we developed a web application using the Shiny framework. This graphical user interface offers all the functionality of the pipeline with a streamlined workflow and can be run locally following installation from https://github.com/gimelbrantlab/magic.

**Figure 1.**
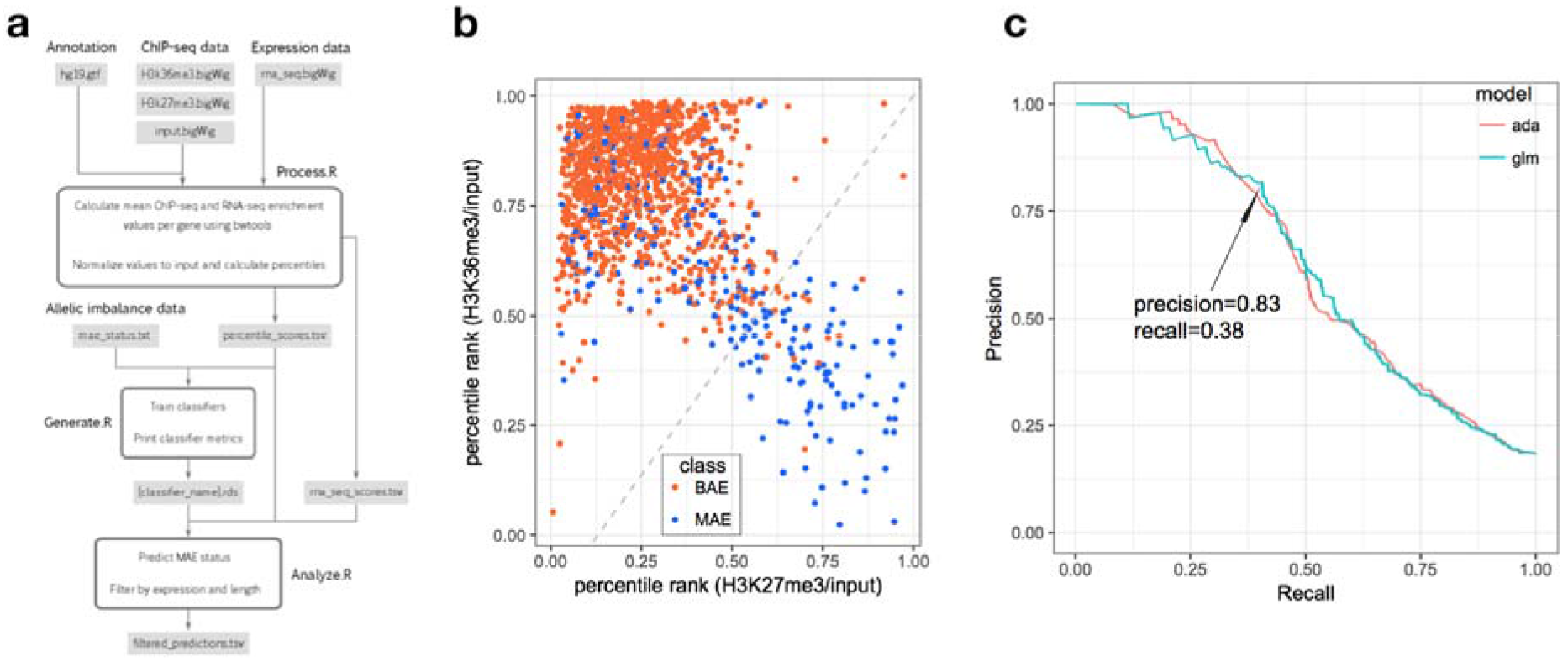
The MaGIC 2.0 pipeline and evaluation of glm performance. **(a)** *Process.R* calculates ChIP-seq enrichment per gene from ChIP-seq and control bigWig files. *Generate.R* trains classifiers using ChIP-seq enrichment and true MAE/BAE calls. *Analyze.R*, uses classifiers to predict MAE/BAE gene status from ChIP-seq data. **(b)** Chromatin signature space of classifier features (H3K27me3 vs. H3K36me3) with the glm’s decision boundary plotted (*dashed line*) with MAE status of true labeled testing data (MAE: *blue*, BAE: *red*). **(c)** Precision-recall curve of glm and ada models evaluated on additional testing data.

## Results

### Software validation

In order to validate MaGIC software, we trained a monoallelic expression classifier using the same datasets as in our previous studies (Nag *et al.*, 2015, 2013). The datasets include ChIP-seq H3K27me3 and H3K36me3 enrichment data for the GM12878 human B-lymphoblastoid cell line (ENCODE Project Consortium, 2012) and a list of 263 monoallelically expressed and 1024 biallelically expressed genes identified in human B-lymphoblastoid clonal cell lines (Gimelbrant *et al.*, 2007).

Because MAE genes make up a small proportion of expressed genes, these datasets are naturally imbalanced. In order to avoid excessive numbers of false positive calls due to this imbalance, we trained the models with optimization parameters for precision and area-under-the-curve of precision-recall (PR) curves, rather than using traditional receiver operating characteristic curves (ROC curves) (Saito and Rehmsmeier, 2015).

We trained a total of 9 models, including a neural network with 2 hidden layers, a support vector machine with multiple kernels, a multi-layer perceptron, three tree-based models (an adaptive boosted classification tree - ada, tree models from genetic algorithms, and recursive partitioning and regression trees), random forest, a K-Nearest Neighbor model and a generalized linear model with stepwise feature selection (glm). After training, all models were tested on an additional human dataset, with 253 MAE genes and 1127 BAE genes identified in monoclonal cell lines derived from GM12878 (Nag *et al.*, 2013) (Dataset S2).

Among all models tested, the generalized linear model had the highest precision value (**Figure 1a, b**; **Table S1**). Depending on the purpose of the analysis, the choice between models can be made using either the precision or the F1 score. The precision score is superior if the user wants to have the lowest number of false positives possible e.g., in identifying high-confidence MAE genes. The F1 score is a balanced metric that is useful if the user is performing genome-wide analysis and wants a higher coverage of MAE genes. However, it should be noted that high precision values come at the expense of “recall” or general coverage of the dataset, as the classifier misses a significant portion of MAE genes in the sample via false negative predictions.

The generalized linear model is packaged along with MaGIC software. It was also evaluated on mouse B-lymphoid clonal cell lines, mouse fibroblasts and mouse neural progenitor cells (data from (Nag et al. 2015), Tables S2 and S3); performed similarly to the classifier from (Nag *et al.*, 2015) (**Table S2**).

### Conclusion

The MaGIC open source toolset described here can be used to map monoallelic expression in a variety of human and mouse cell types using existing models. It can also employed to train new models with a different set of chromatin marks or genomic features, while assisting the user in identifying the best suited algorithm. As epigenomic data is becoming increasingly available in many cell and tissue types, we believe the versatility of the MaGIC pipeline will prove invaluable to investigate MAE mechanisms, function, and contribution to disease.

## Acknowledgements

Funding: R01 GM114864 to AAG; SS and HW were students in a program supported by NIH award U54 HG007963. No conflict of interest declared.

## References

Chess,A. (2016) Monoallelic Gene Expression in Mammals. Annu. Rev. Genet., 50, 317–327.

ENCODE Project Consortium (2012) An integrated encyclopedia of DNA elements in the human genome. Nature, 489, 57–74.

Gimelbrant,A. et al. (2007) Widespread monoallelic expression on human autosomes. Science, 318, 1136–1140.

Kent,W.J. et al. (2010) BigWig and BigBed: enabling browsing of large distributed datasets. Bioinformatics, 26, 2204–2207.

Kuhn,M. (2008) Building Predictive Models inRUsing thecaretPackage. J. Stat. Softw., 28.

Nag,A. et al. (2015) Chromatin Signature Identifies Monoallelic Gene Expression Across Mammalian Cell Types. G3, 5, 1713–1720.

Nag,A. et al. (2013) Chromatin signature of widespread monoallelic expression. Elife, 2, e01256.

Pohl,A. and Beato,M. (2014) bwtool: a tool for bigWig files. Bioinformatics, 30, 1618–1619.

Saito,T. and Rehmsmeier,M. (2015) The precision-recall plot is more informative than the ROC plot when evaluating binary classifiers on imbalanced datasets. PLoS One, 10, e0118432.

Savova,V. et al. (2013) Autosomal monoallelic expression: genetics of epigenetic diversity? Curr. Opin. Genet. Dev., 23, 642–648.

Savova,V. et al. (2016) Genes with monoallelic expression contribute disproportionately to genetic diversity in humans. Nat. Genet., 48, 231–237.

Savova,V. et al. (2017) Risk alleles of genes with monoallelic expression are enriched in gain-of-function variants and depleted in loss-of-function variants for neurodevelopmental disorders. Mol. Psychiatry, 22, 1785–1794.

